# Tumor cells upregulate *CCL22* in a STING-dependent manner in response to paracrine factors released by STING-activated myeloid cells and type I interferons

**DOI:** 10.1101/2025.07.08.663704

**Authors:** Elmira M Lomashvili, Jihyun Kim, Lingwei Kong, Pamela R. Cook

## Abstract

Immunosuppressive elements within the tumor microenvironment include both regulatory T cells (Tregs) and M2 macrophages. A well-described mechanism of Treg recruitment occurs via the chemokine CCL22, and CCL22 has also recently been implicated in the polarization of tumor-associated macrophages to the M2a subtype. Our lab and others have shown that CCL22 is upregulated in cancer cells following activation of the <u>ST</u>imulator of <u>IN</u>terferon <u>G</u>enes (STING). STING triggers immune responses against pathogenic and self-DNA mislocalized to the cytoplasm, which can accumulate in cancer cells due to chromosomal instability, damaged mitochondria, and increased expression of LINE-1 retrotransposons. STING activation has been associated with both anti- and pro-tumor immune responses, and a potential mechanism of STING-mediated immune evasion is through CCL22 upregulation. CCL22 was first characterized in macrophages, and here we investigate the effects of STING activation on *CCL22* expression in macrophages and monocytes. We report that human macrophages and monocytes are resistant to *CCL22* upregulation by STING, but that STING-activated macrophages and monocytes release unidentified paracrine factor(s) that dramatically increase *CCL22* upregulation in cancer cells in a manner that remains STING-dependent, as evidenced by the inability of STING knockout cells to upregulate *CCL22* in response to these factors. We further found that exogenous type I interferons (IFNs), a major downstream product of STING activation, also upregulate *CCL22* in cancer cells via a STING-dependent mechanism and that exogenous IFNβ can directly activate STING.

## INTRODUCTION

The <u>ST</u>imulator of <u>IN</u>terferon <u>G</u>enes (STING) is an effector protein that transmits signals from various cytosolic DNA sensors to transcription factors, including interferon regulatory factor 3 (IRF3), which leads to the expression of type I interferon (*IFN*) genes and downstream immune responses (1–3). Healthy, non-mitotic eukaryotic cells confine DNA to the nucleus and mitochondria; thus, the presence of cytosolic DNA often signals infection by microbes, making STING an important sentinel in innate immune surveillance. Cancer cells are often characterized by self-DNA mislocalized to the cytoplasm due to chromosomal instability, damaged mitochondria, and reactivated LINE-1 retrotransposons (4, 5). LINE-1-mediated activation of STING is thought to contribute to inflammation in multiple disease states, including age-related neuropathologies and cancer (5–8). In the context of cancer, LINE-1 is highly expressed in many epithelial tumors (9–11), and its activation of STING in those cells may have important clinical implications on host anti-tumor immune responses. STING activation in cancer can boost anti-tumor immunity due to the release of type I IFNs, and indeed STING agonists represent promising immunotherapy agents (12, 13). However, the roles of STING in carcinogenesis, tumor progression, and metastasis are complex, and in some cases STING activation has been associated with impaired anti-tumor immunity (14–16).

Our lab and others have demonstrated a link between STING activation and increased expression of a chemokine associated with immunosuppression in the tumor microenvironment, the C-C-motif ligand 22 (*CCL22*) (17–19). Correlations between *CCL22* expression and adverse outcomes across various cancer types have been reported in multiple studies (18, 20–24), although CCL22 has also been associated with increased patient survival in some cancers, particularly for colon and head and neck squamous cell carcinomas (19 and references therein). CCL22 recruits regulatory T-cells (Tregs) via its interaction with the CCR4 receptor, which may reduce tumor immune responses and promote immunosuppression in some cancers (25, 26). Recently, CCL22 has also been reported to have a role in polarizing tumor-associated macrophages to the M2a pro-tumor subtype (27).

CCL22, also known as macrophage-derived chemokine (MDC), was originally discovered in macrophages (28, 29). Macrophages and other myeloid cells are major targets of STING agonists in the tumor microenvironment (27). Here we sought to investigate the relationship between STING activation and *CCL22* upregulation in macrophages and monocytes. Although we found that STING activation did not increase *CCL22* expression in macrophages or monocytes, we found that factor(s) released from these myeloid cells dramatically upregulated *CCL22* in cancer cells in a paracrine fashion. Moreover, we determined that this paracrine effect remains dependent on STING in the cancer cells, based on results from our STING knockout cell line. We previously showed that *CCL22* upregulation in response to STING activation occurs predominantly via the STING-IRF3 axis (19), the same axis that triggers robust upregulation of type I IFNs. We thus hypothesized that type I IFNs could be the relevant paracrine factor(s) released by myeloid cells that upregulate *CCL22* in cancer cells. Although we found that exogenous type I IFNs did indeed increase *CCL22* expression in cancer cells, the myeloid-derived paracrine factors tended to have a larger effect, particularly in HeLa cells. Surprisingly, we also found that upregulation of *CCL22* in response to type I IFNs remained dependent on STING, and that IFNβ itself could activate STING.

## MATERIALS AND METHODS

### Cells and cell culture

Cell lines used in the study include MCF7 (human mammary gland adenocarcinoma), THP-1 (human monocytic leukemia), both purchased from ATCC, human HeLa cervical adenocarcinoma cells, a kind gift from Dr. Anthony Furano from the National Institute of Diabetes and Digestive and Kidney Diseases as described in (19), and HeLa^-/-*TMEM173*^ STING *(TMEM173)* knockout cells, engineered from the aforementioned wild type HeLa with the CRISPR-Cas 9 system, also described in (19). All cells were cultured in DMEM high glucose with GlutaMAX and pyruvate (Gibco, cat. 10569010) supplemented with 10% fetal bovine serum (Gibco OneShot, non-heat-inactivated, cat. A3160401) and 1x antibiotic-antimycotic (100 units/mL penicillin, 100 ug/mL streptomycin, 0.25 ug/mL Amphotericin B; Gibco, cat. 15240096) in a humidified incubator at 37°C with 5% CO2. THP-1 culture media was additionally supplemented with 1x 2-mercaptoethanol (Gibco, cat. 21985023).

### Cell culture treatments (STING agonists, IFNα, IFNβ, and MMHAR-2 block)

Four commercially available STING agonists were used in the study: a stabilized analog of the cyclic dinucleotide canonical STING agonist 2’3’-cGAMP (2’3’-cGAM(PS)2(Rp/Sp)), hereafter referred to as cGAM(PS)2 (Invivogen, cat. tlrl-nacga2srs), a stabilized cyclic adenine monophosphate-inosine monophosphate (cAIM(PS)2 Difluor (Rp/Sp), hereinafter referred to as cAIM(PS)2 (Invivogen, cat. tlrl-nacairs-2), E7766, a macrocycle-bridged diammonium salt (Chemietek, cat. CT-E7766), and the diamidobenzimidazole diABZI (Selleckchem, cat. S8796). All agonists except diABZI were water soluble and reconstituted using a companion vial of sterile, endotoxin-free LAL water; diABZI was reconstituted in dimethyl sulfoxide (DMSO). All agonists were diluted to a final concentration of 30 uM in complete cell culture media and applied to cells as specified in the figures. Human recombinant IFNα stock (pbl assay science, cat. 11101-1) was diluted to concentrations indicated in the figure legends in complete cell culture media. Lyophilized recombinant human IFNb (PeproTech, cat. 300-02) was initially reconstituted in sterile, endotoxin-free LAL water and further diluted into sterile 0.1% bovine serum albumin (BSA) in 1x phosphate-buffered saline (PBS) to 100 ng/uL and further diluted to concentrations indicated in the figure legends in complete cell culture media before adding to cells. MMHAR-2 blockade was performed by pretreating cells with 8.3 μg/mL of the MMHAR-2 antibody, then adding indicated treatments diluted in cell culture media, reducing the final concentration of MMHAR-2 to 5 μg/mL. Cells were then incubated for approximately 30 hours prior to harvesting for either Western blots or RTqPCR.

### Macrophage differentiation and generation of conditioned media

THP-1 cells were differentiated into macrophages by plating cells in complete culture media supplemented with 150 nM phorbol 12-myristate 13-acetate (PMA) and incubating for 96 hours, with one media change performed after 48 hours to replenish PMA. PMA-containing media was then removed, and cells were allowed to rest for 24 hours prior to treatment with cGAM(PS)2. Conditioned media from both macrophages and undifferentiated monocytes was obtained by treating cells with fresh complete media, with or without the addition of 30 uM cGAM(PS)2, incubating for 24 hours, then gently removing media, centrifuging at 300 x g, and transferring to cancer cells.

### Cell imaging

The morphological changes during THP-1 differentiation were imaged using an EVOS FLoid system.

### RNA purification and RTqPCR

Cells were harvested using TRIzol reagent (Invitrogen, cat. 15596026) followed by column RNA purification and on-column DNase digestion according to the TRIzol two-step protocol from the Monarch total RNA miniprep kit (NEB, cat. T2010S). Purity and concentration of the RNA were assessed using spectrophotometry. First-strand cDNA synthesis was performed using the LunaScript RT SuperMix Kit (NEB, cat. E3010S) or LunaScript (-) RT SuperMix control with added M-MuLV Reverse Transcriptase (NEB). qPCR was performed on a QuantStudio 3 real-time PCR machine using TaqMan Fast Advanced Master Mix (Applied Biosystems, cat. A44359). Singleplexed and duplexed 5’ endonuclease assays were performed using the following TaqMan Assays: *CCL22* (Hs00171080); IFN-β (Hs01077958_s1; beta actin (Hs99999903); and 18s (Hs99999901). For duplexed assays, a beta-actin primer-limited assay (IDT) was used. Samples for qPCR were assayed in technical triplicates, and data were analyzed with the delta-delta Ct method; when multiple housekeeping genes were measured, their combined geometric mean was used for analysis.

### Cell lysis and Western blotting

Cells were lysed for immunoblotting in 3% SDS, 25 mM Tris-HCl pH 7.4, and 0.5 mM EDTA supplemented with 3X HALT protease and phosphatase inhibitor cocktail (ThermoFisher, cat. 78440). Lysate homogenization was achieved with QIAshredder columns (Qiagen, cat. 79656), after which protein concentration was determined using the BioRad DC Protein Assay (cat. 5000112). Equal amounts of lysates were resolved with SDS-PAGE. Transfers were performed using a wet tank system overnight at 28 V or the BioRad Trans-Blot Turbo Transfer system. Blots were imaged with an infrared Li-Cor Odyssey CLx Imager and processed using Image Studio (Li-Cor). Positive controls for phospho-STING (S366) were prepared as commercially described by transfecting THP-1 differentiated macrophages with dsDNA (pcDNA3.1 (+) puro), (3.33 ug/mL) using Opti-MEM (Gibco, cat. 31-985-062) as the DNA diluent and TransIT-LT1 (Mirus, cat. MIR 2300) at a ratio of 1 ug DNA: 2 uL TransIT-LT1; cells were harvested 4 hours after transfection with the lysis buffer described above.

### Antibodies

Antibodies for Western blots included: anti-phospho(S366)-STING (Cell Signaling Technology, cat. 19781S); anti-phospho(Y701)-Stat1 (Cell Signaling Technology, cat. 9167); anti-IFNAR2 (Cell Signaling Technology, cat. 53883); and anti-beta-tubulin (Abcam, cat. ab6046). The antibody used for IFNAR blockade was anti-human interferon alpha/beta receptor chain 2, clone MMHAR-2 (pbl assay science, cat. 21385-1).

### Statistical analyses

Statistical analyses were performed using GraphPad Prism 9. Tests for significance are described in the figure legends, and statistical significance was defined as p-value ≤ 0.05.

## RESULTS

### STING-activated macrophages and monocytes do not upregulate *CCL22*

Macrophages are a major source of CCL22, reflected in the historical name of CCL22, macrophage-derived chemokine (MDC). Since one of the targets of STING agonists are tumor-associated macrophages, we sought to determine the effect of STING activation on *CCL22* in these cells. Using the well-established THP-1 monocyte-to-macrophage differentiation model (Fig. 1A), we incubated THP-1 monocytes with phorbol myristate acetate (PMA) for four days to establish adherent, differentiated macrophages. Following differentiation, we treated the THP-1-derived macrophages with the canonical human STING agonist cGAM(PS)2 for 24 hours. Unexpectedly, we found that cGAM(PS)2 did not increase *CCL22* expression but rather reduced it (Fig. 1B). To ensure that we had not missed an increase in *CCL22* at an earlier time point, we monitored *CCL22* expression at 0, 6, 12, 24, and 48 hours, which revealed no significant increase at any time point (Fig. S1). Rather, the time course in Fig. S1 shows a downward trend in *CCL22* expression, although these decreases did not reach statistical significance in the context of a Kruskal-Wallis multiple comparisons test, which generally has less power to identify differences in means compared with the *t* test used to analyze data in Figure 1B. Indeed, when analyzing only the 24-hour time-course data with an unpaired, two-tailed *t* test, the decrease in *CCL22* was again significant (p = 0.0290). To determine whether failure to activate *CCL22* in macrophages was due to a lack of STING activation, we confirmed that cGAM(PS)2 induced STING phosphorylation in macrophages at both 6 and 24 hour time points (Fig. 1C). Consistent with results from macrophages, undifferentiated THP-1 monocytes also failed to upregulate *CCL22* in response to cGAM(PS)2 (Fig. 1D).

**Figure 1.**
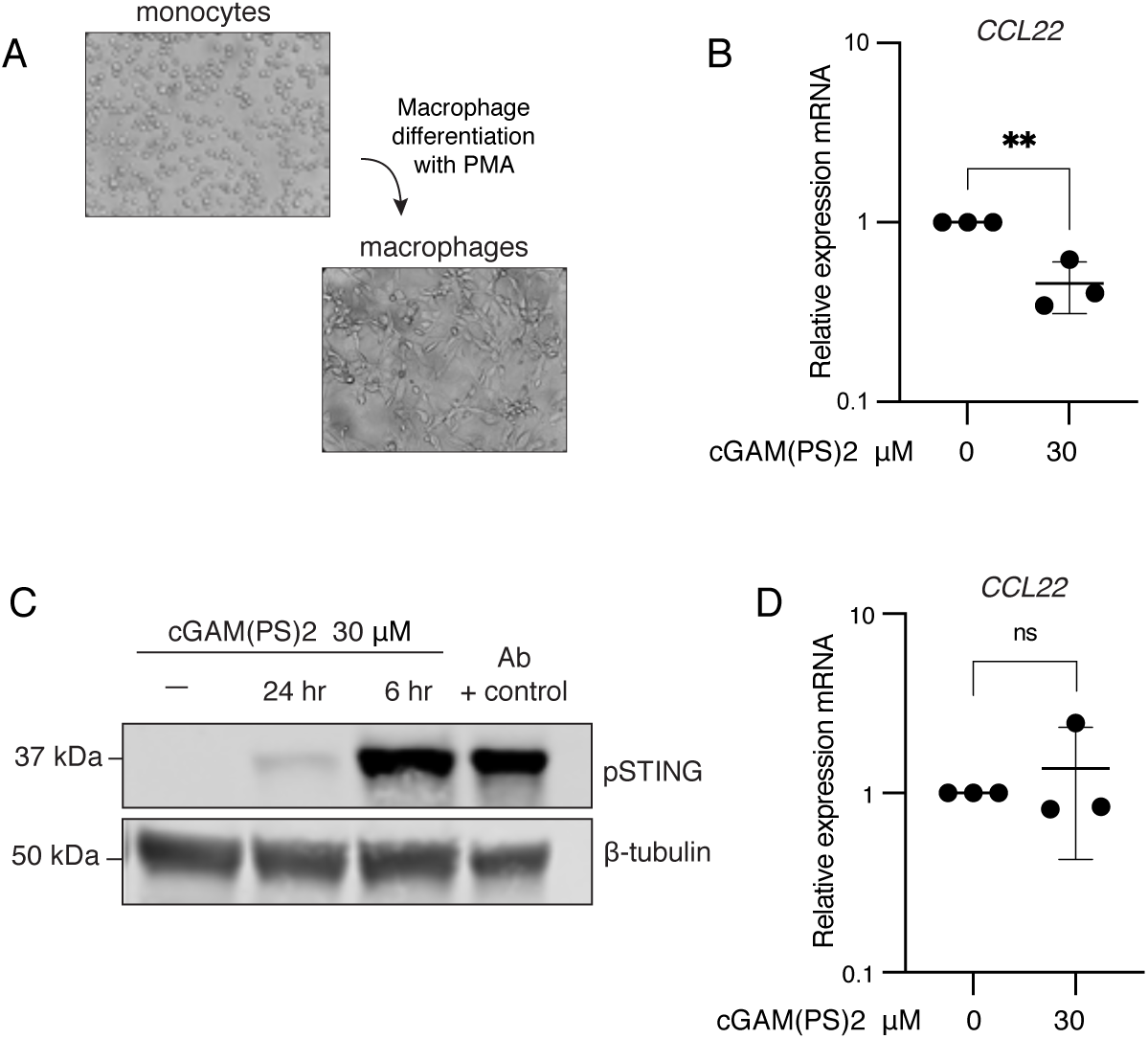
STING-activated macrophages and monocytes do not upregulate *CCL22*. ***A)*** Microscopic images (20x brightfield) of THP-1 monocytes (upper left) and fully differentiated THP-1-derived macrophages (lower right) are shown. ***B)*** THP-1-derived macrophages were treated with cGAM(PS)2 (30 μM) for 24 hours, then harvested in TRIzol for RNA purification and RTqPCR. Relative *CCL22* gene expression is shown. ***C)*** THP-1-derived macrophages were treated with cGAM(PS)2 (30 μM) for 6 and 24 hours, then whole cell lysates (20 μg) and antibody positive control (7 μg) were resolved with SDS-PAGE, transferred to nitrocellulose, and probed for phosphorylated STING and beta-tubulin. ***D)*** THP-1 monocytes were treated with cGAM(PS)2 (30 μM) and harvested for RTqPCR and *CCL22* gene expression as described in *B*. Error bars represent standard deviations of independent biological replicates. Statistical analyses were performed with unpaired, two-tailed *t* test (*B)* or non-parametric Wilcoxon signed-rank test (*D*) due to non-Gaussian distribution of the treated sample per Shapiro-Wilk test.

### STING-activated macrophages and monocytes release factor(s) that upregulate *CCL22* in cancer cells

Macrophages have important roles in anti-tumor immunity but can also contribute to immunosuppressive environments (30). We thus sought to determine whether STING-activated macrophages might release factors that could act via paracrine signaling to alter *CCL22* expression in cancer cells. *CCL22* has been reported to play a role in breast (20, 22, 24) and cervical (18, 23) cancers, and we used human breast (MCF7) and cervical (HeLa) cancer cells in these experiments. These cell lines were also used in our previous study, where we showed that both CCL22 mRNA and protein were increased upon direct STING activation with cytosolic DNA or cGAM(PS)2 (19). To investigate potential paracrine activation of *CCL22* in these cells, we first treated macrophages with cGAM(PS)2 or a mock control for 24 hours, then removed the media and applied it to cancer cells for 24 hours. Both MCF7 (Fig. 2A) and HeLa cells (Fig. 2B) exposed to conditioned media from macrophages treated with cGAM(PS)2 exhibited a robust increase in *CCL22* compared to media from untreated macrophages or treatment with cGAM(PS)2 alone, although conditioned media from untreated macrophages also increased *CCL22* in HeLa cells, albeit to a lower extent. Monocytes also released factors that increased *CCL22* above levels induced by direct STING activation in MCF7 (Fig. 2C) and HeLa (Fig. 2D) cells.

**Figure 2.**
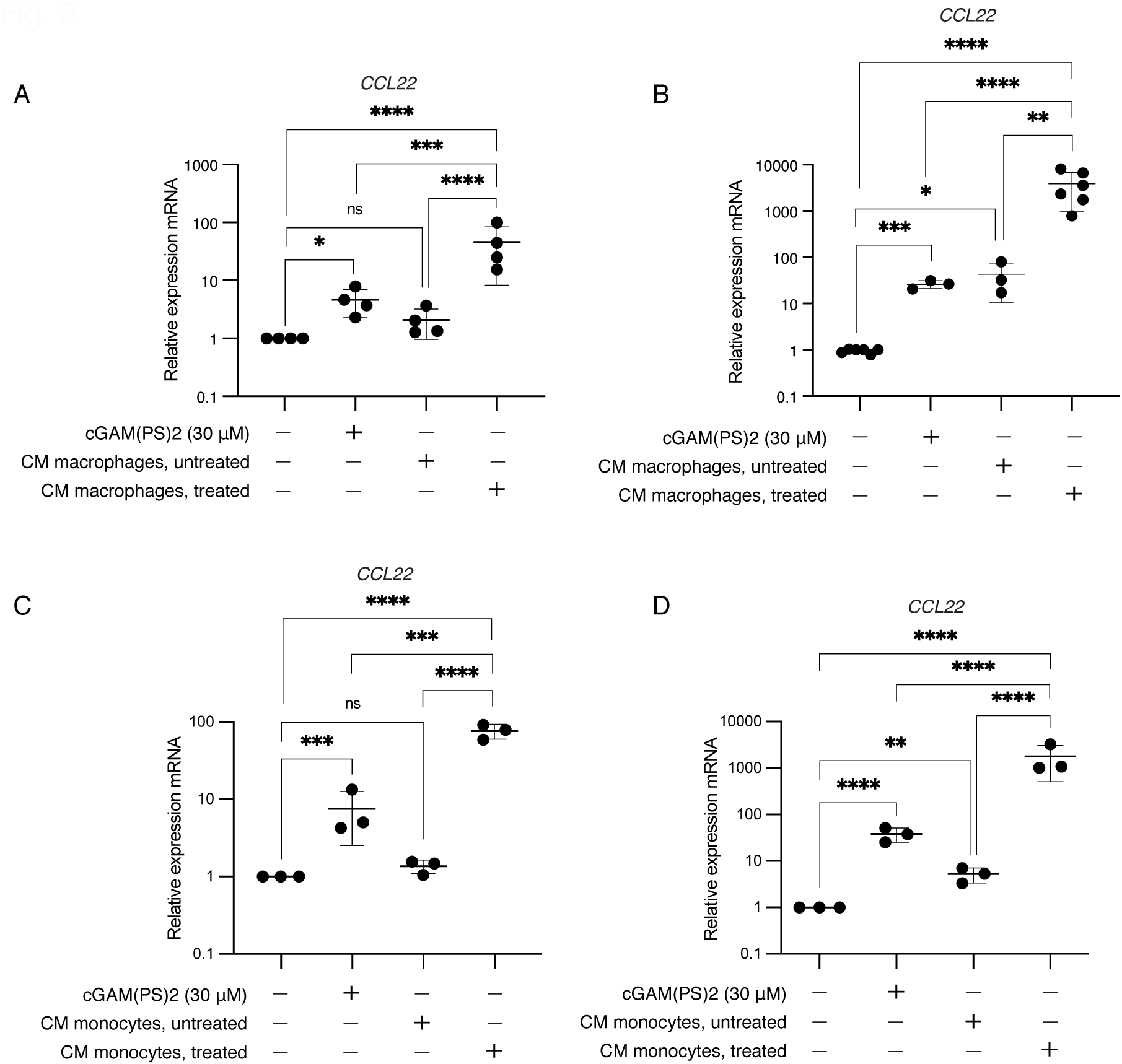
STING-activated macrophages and monocytes release factor(s) that upregulate *CCL22* in cancer cells. ***A)*** MCF7 and ***B)*** HeLa cells were treated for 24 hours with cGAM(PS)2 (30 μM) or conditioned media (CM) from macrophages that were either untreated or treated with cGAM(PS)2 (30 μM) for 24 hours prior to media collection. Cells were harvested with TRIzol for RNA purification and RTqPCR to assess relative expression of *CCL22. **C)*** MCF7 and ***D)*** HeLa cells were treated for 24 hours with cGAM(PS)2 (30 μM) or conditioned media (CM) from monocytes that were either untreated or treated with cGAM(PS)2 (30 μM) for 24 hours prior to media collection. Cells were harvested with TRIzol for RNA purification and RTqPCR to determine expression of *CCL22* as above. For all figures, error bars represent standard deviations of independent biological replicates. Statistical analyses were performed with one-way ANOVAs on log-transformed data and Tukey’s pairwise comparisons (*A*, *C*, and *D*) or with Brown-Forsythe and Welch ANOVA (*B*) due to significant differences among standard deviations.

### Paracrine activation of *CCL22* in cancer cells activates STING and is dependent on STING

Our previous study showed that *CCL22* upregulation in response to cytosolic dsDNA and cGAM(PS)2 in HeLa cells was dependent on STING. However, in the current experiments, it was possible that paracrine-mediated activation of *CCL22* in these cells was independent of STING. To investigate this, we first determined whether conditioned media from STING-activated macrophages had any effect on STING activation in HeLa cells, as determined by analysis of STING phosphorylation on serine 366 (31, 32). Figure 3A shows that STING in HeLa cells was indeed activated by media obtained from STING-activated macrophages, and this activation was greater than when HeLa cells were treated directly with cGAM(PS)2. Importantly, in our previous study, we confirmed that free-floating DNA, which could possibly have been present in the conditioned media, did not activate STING in HeLa cells (19). Notably, the degree of STING activation by each treatment in Figure 3A generally corresponds to relative levels of *CCL22* upregulation in Figure 2B, including the slight increases observed in response to media from untreated macrophages. Similar results for STING phosphorylation were observed when treating HeLa cells with conditioned media from monocytes (Fig. S2A), which again parallels increases in *CCL22* in Figure 2D. Note that we did not examine STING phosphorylation on S366 in MCF7 cells due to our previous finding that STING activation in MCF7 cells with cytosolic dsDNA or pharmacological agonists occurred in the absence of phosphorylation on S366 for unknown reasons, which are further discussed in that study (19). To determine whether the observed STING activation was *required* for paracrine-mediated upregulation of *CCL22*, we used a STING (gene name *TMEM173*) knockout cell line, HeLa^-/-^ *^TMEM173^*, which we previously established and characterized from the parental HeLa cells used here (19). Treatment of the HeLa^-/-*TMEM173*^ STING knockout cells with media from STING-activated macrophages showed no statistically significant increase in *CCL22* expression, indicating that intact STING signaling in the cancer cells was still required for upregulated *CCL22* (Fig. 3B). Similar results were obtained using media from STING-activated monocytes (Fig. S2B).

**Figure 3.**
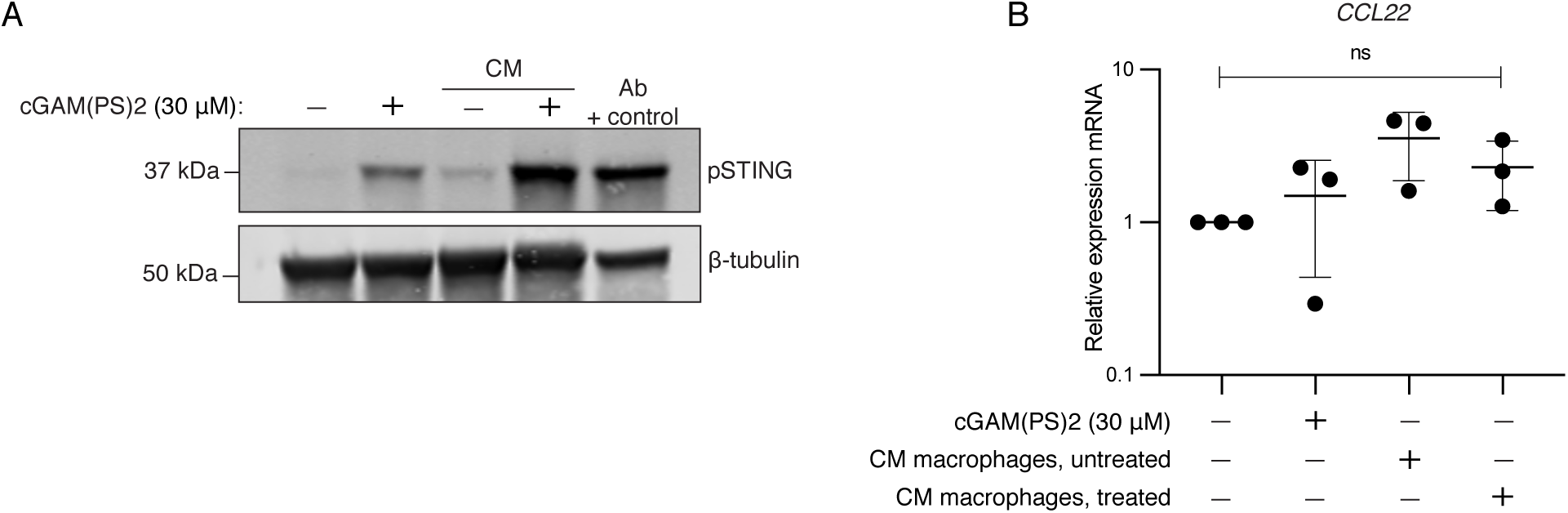
Paracrine activation of *CCL22* in cancer cells activates STING and is dependent on STING. ***A)*** HeLa cells were treated for 24 hours with cGAM(PS)2 (30 μM) or conditioned media (CM) from macrophages that were either untreated or treated with cGAM(PS)2 (30 μM) for 24 hours. Whole cell lysates (25 μg) and antibody positive control (5 μg) were resolved with SDS-PAGE, transferred to nitrocellulose and probed for phosphorylated STING and beta-tubulin. The image shown is representative of at least three independent experiments. ***B)*** HeLa^-/-^*^TMEM173^* STING knockout cells were treated for 24 hours with cGAM(PS)2 (30 μM) or conditioned media (CM) from macrophages that were either untreated or treated with cGAM(PS)2 (30 μM) for 24 hours prior to media collection. Cells were harvested with TRIzol for RNA purification and RTqPCR to determine relative expression of *CCL22*. Error bars represent standard deviations of independent biological replicates. Statistical analysis was performed using one-way ANOVA and Tukey’s pairwise comparisons.

### Type I IFNs have modest effects on *CCL22* in HeLa cells that remain STING-dependent

We previously found that STING-mediated *CCL22* upregulation in HeLa cells is mediated predominantly via the STING-IRF3 axis, which triggers robust expression of type I IFNs (19). We therefore hypothesized that the relevant paracrine factor(s) leading to the increase in *CCL22* might be type I IFNs. Overall however, IFNα failed to elicit strong *CCL22* expression in either cell type, and effects of IFNβ plateaued in dose-response curves, indicating that the relevant paracrine factor was unlikely a higher concentration of type I IFNs.

In HeLa cells, IFNβ did not significantly upregulate *CCL22* at any dose at 24 hours (Fig. S3A), and subsequent time-course analyses indicated the maximum response to both IFNα and IFNβ occurred at 48 hours. However, even at 48 hours, IFNα failed to significantly upregulate *CCL22* in HeLa cells except at the highest dose (100 ng/mL), and then only to an average of 15 fold (Fig. 4A). IFNβ increased *CCL22* at the lowest dose (1 ng/mL) in HeLa cells, but only to approximately 10 fold, and higher doses of IFNβ failed to elicit any additional increase (Fig. 4B). MCF7 cells upregulated CCL22 in response to type I IFNs more rapidly than HeLa cells, showing higher levels at 24 hours compared to 48 hours (Fig. S3B). At the highest dose, IFNα increased *CCL22* by an average of only 6.7 fold in 24 hours (Fig. 4C). Paralleling the relative effects of IFNα and IFNβ in HeLa cells, IFNβ also increased *CCL22* in MCF7 cells more than IFNα, with an average of 57 fold change in 24 hours at the highest dose (Fig. 4D). However, the effect of IFNβ was still below that observed from paracrine activation of MCF7 cells by conditioned media from STING-activated monocytes (76.5 fold, Fig. 2C), though it was slightly above levels induced by conditioned media from STING-activated macrophages (46.3 fold, Fig. 2A).

**Figure 4.**
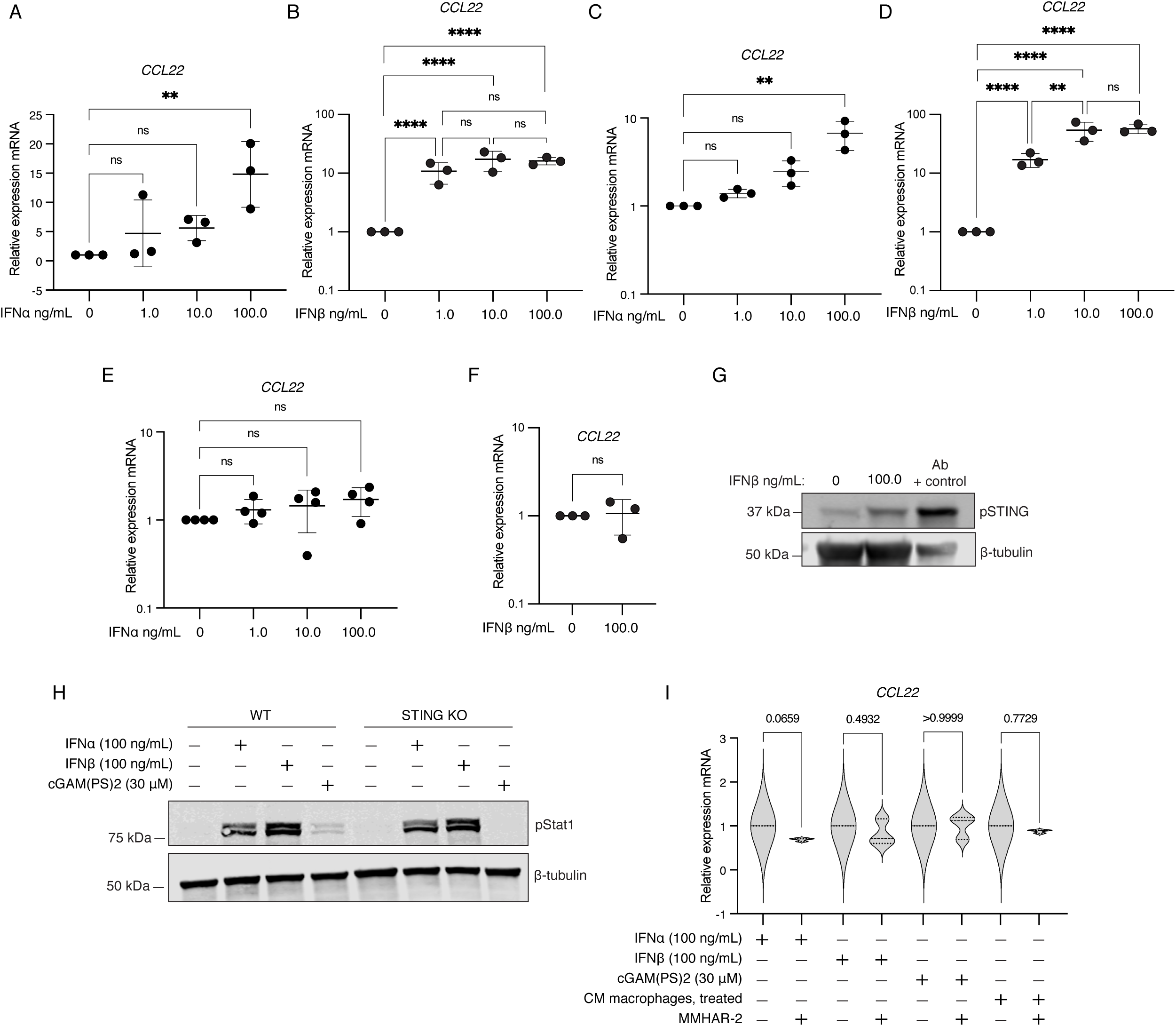
Type I IFNs have modest effects on *CCL22* in HeLa cells that remain STING-dependent. HeLa ***(A-B)*** and MCF7 cells ***(C-D)*** were treated with exogenous type I IFNs at indicated concentrations for 48 (HeLa) or 24 hours (MCF7), then harvested with TRIzol for RNA purification and RTqPCR to determine relative gene expression of *CCL22*. ***E-F)*** HeLa ^-/-^*^TMEM173^* STING knockout cells were treated with exogenous type I IFNs at indicated concentrations for 48 hours, then harvested with TRIzol for RNA purification and RTqPCR analysis of *CCL22.* Error bars represent standard deviations of independent biological replicates. Statistical analyses were performed using one-way ANOVA with Dunnett’s multiple comparisons test (*A, C, E*), one-way ANOVA on log-transformed data with Tukey’s pairwise comparisons (*B, D*), and unpaired, two-tailed t test (*F*). ***G)*** HeLa wild type cells were treated with IFNβ (100 ng/mL) for 48 hours, and whole cell lysates (60 μg) and antibody positive control (5 μg) were resolved with SDS-PAGE, transferred to nitrocellulose and probed for phosphorylated STING and beta-tubulin. ***H)*** HeLa wild type (WT) and HeLa ^-/-*TMEM173*^ STING knockout (KO) cells were treated as indicated for approximately 30 hours prior to harvesting; whole cell lysates (15 μg) were resolved with SDS-PAGE, transferred to nitrocellulose, and probed for phosphorylated Stat1 and beta-tubulin. ***I)*** HeLa cells were pretreated for 3 hours with MMHAR-2 neutralizing antibody, then indicated treatments were added, including conditioned media (CM) from macrophages treated with cGAM(PS)2 (30 μM), and cells were incubated for approximately 48 hours prior to harvest in TRIzol for RNA purification and RTqPCR to assess relative changes in *CCL22* expression. Statistical analyses were performed using one-way ANOVA with Šídák’s multiple comparisons test.

We next tested the effect of exogenous IFNα and IFNβ on *CCL22* upregulation in the context of STING knockout cells, HeLa^-/-*TMEM173*^. Neither IFNα (Fig. 4E) nor IFNβ (Fig. 4F) significantly affected *CCL22* expression in STING knockout cells. These data were unexpected and indicate that the effect of exogenous IFNα and IFNβ on *CCL22* is dependent on STING. Although STING is well-known to trigger expression of type I *IFNs* downstream of IRF3, to our knowledge, a requirement of STING for the effects of exogenous type I IFNs has not been reported. To determine whether STING was activated by type I IFN signaling, we treated wild-type HeLa cells with IFNβ and then performed a Western blot to assess STING phosphorylation, and indeed found that exogenous IFNβ resulted in activated STING (Fig. 4G). We did not test the effect of IFNα due to the comparatively minimal effect we observed from IFNα on *CCL22* relative to IFNβ. To rule out the possibility that type I IFNs failed to upregulate *CCL22* in the STING KO cells due to an inadvertent disruption of the interferon receptor (IFNAR) and downstream Stat1 signaling, we confirmed that our STING KO cells exhibited the same degree of Stat1 phosphorylation in response to IFNα and IFNβ as wild-type cells (Fig. 4H). As expected, we found that cGAM(PS)2, which activates STING and STING-mediated type I IFNs, also slightly induced Stat1 in the wild-type cells, but this activation was blocked in the STING KO cells, reflecting the lack of STING-mediated upregulation of type I IFNs available to bind and activate the IFNAR-Stat1 pathway in those cells (Fig. 4H). Confirming intact pStat1 signaling in the STING KO cells is consistent with our hypothesis that *CCL22* upregulation in response to exogenous type I IFN is mediated through STING, not Stat1. Given that Stat1 phosphorylation was equivalently induced in wild-type and STING knockout cells by type I IFNs, it was also unlikely that IFNAR expression would have been reduced in the STING knockout cells, but to confirm this, we also show equivalent levels of IFNAR in both cell lines (S3C).

The inability of type I IFNs to activate *CCL22* expression in our STING KO cells, combined with confirmation of intact IFNAR expression and signaling in STING KO cells, indicated that *CCL22* upregulation in wild-type cells by type I IFNs occurs via STING. To further examine the role of IFNAR signaling on *CCL22* activation in the context of type I IFNs, direct STING activation with cGAM(PS)2, or paracrine activation using conditioned media, we performed an IFNAR blockade using a well-characterized neutralizing monoclonal antibody from PBL Assay Science, anti-human IFN-alpha/beta receptor chain 2, clone MMHAR-2. To first confirm that MMHAR-2 exerted a neutralizing effect on IFNAR signaling, we analyzed the ability of this antibody to reduce Stat1 phosphorylation in response to IFNα and IFNβ, as well as cGAM(PS)2. MMHAR-2 somewhat reduced levels of phosphorylated Stat1 in response to IFNα, but not IFNβ (Fig. S3D). This difference might be attributable to the reported higher affinity of IFNβ for IFNARs compared to IFNα (33, 34). Given that cGAM(PS)2 upregulates type I IFNs through its activation of STING, we also observed as expected that the MMHAR-2 blockade reduced Stat1 phosphorylation following treatment with cGAM(PS)2 (Fig. S3D). We also confirmed that MMHAR-2 exerted no activating effect on Stat1 (Fig. S3E). After establishing the neutralizing potential of MMHAR-2 in our system, we sought to determine the contribution of IFNAR signaling to *CCL22* upregulation in response to type I IFNs, cGAM(PS)2, and paracrine factors from STING-activated macrophages. Figure 4I shows the relative degree of inhibition induced by MMHAR-2 for each treatment. Although not statistically significant, MMHAR-2 produced an observable downward trend in *CCL22* expression for all treatments except cGAM(PS)2. The neutralizing effect of MMHAR-2 was most noticeable in cells treated with IFNα, where the p-value is approaching significance at 0.0659, and these results parallel those observed for MMHAR-2 inhibition of IFNα-mediated Stat1 phosphorylation in Figure S3D. Also in Figure S3D, the very slight increase in Stat1 phosphorylation from cGAM(PS)2 treatment, as well as its return to baseline with MMHAR-2 pretreatment, most likely reflects the activity of cGAM(PS)2-induced type I IFNs downstream of STING. That the IFNAR block had the least effect on *CCL22* upregulation by cGAM(PS)2 is consistent with our other data indicating that *CCL22* upregulation downstream of STING in HeLa cells is not mediated primarily or solely by type I IFNs.

### STING agonists differentially upregulate *CCL22* and *IFNβ*

We were curious, given the differential effects of type I IFNs and cGAM(PS)2 on *CCL22,* whether different STING agonists would have distinctive effects on *CCL22* compared to *IFNβ*. We tested a panel of four STING agonists that included two cyclic and two and non-cyclic dinucleotides. The cyclic dinucleotides included cGAM(PS)2 and the stabilized analog of adenosine-inosine monophosphate (cAIMP(PS)2) (35, 36). The two non-cyclic dinucleotides were diamidobenzimidazole (diABZI) (37–39) and E7766, a locked, macrocycle-bridged agonist (38, 40, 41). In HeLa cells, all agonists increased *CCL22* but had no significant effect on *IFNβ* expression (Fig. 5A-5D). Three of the STING agonists increased *IFNβ* in MCF7 cells to statistically significant levels, but *CCL22* was increased in response to all agonists, including diABZI, the agonist that did not statistically increase *IFNβ* (Fig. 5E-5H). These results are consistent with a mechanism of *CCL22* induction via STING that does not rely exclusively on type I IFNs.

**Figure 5.**
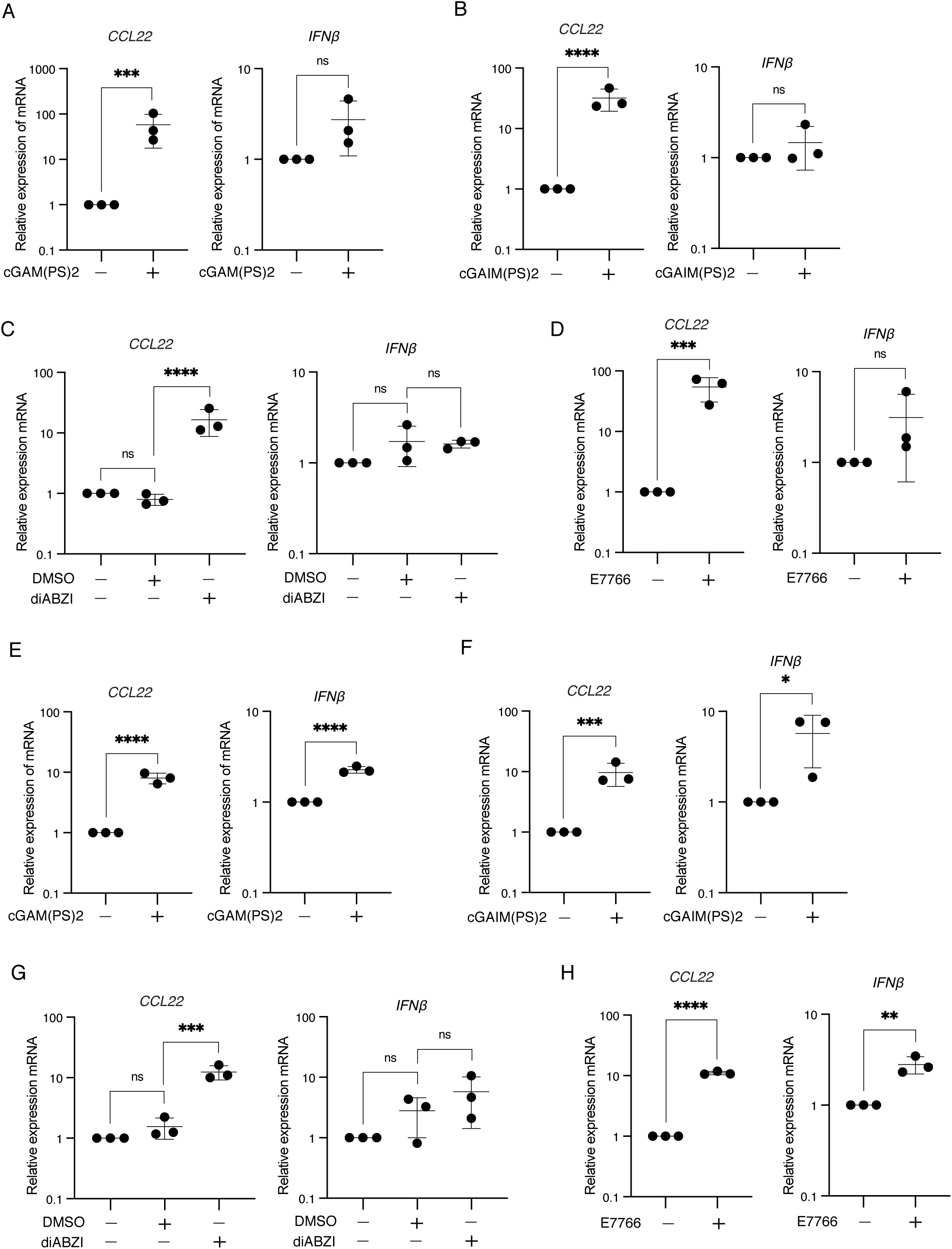
STING agonists differentially upregulate *CCL22* and *IFNβ.* HeLa ***(A-D)*** and MCF7 cells ***(E-H)*** were treated with 30 μM of each indicated STING agonist or DMSO vehicle control (for diABZI) for 24 hours, then harvested in TRIzol for RNA purification and RTqPCR using TaqMan assays for *CCL22* or *IFNb*. Error bars represent standard deviations of independent biological replicates. Statistical analyses were performed on log-transformed data using unpaired, two-tailed t tests except for *C* and *G*, which were analyzed with a one-way ANOVA and Šídák’s multiple comparisons test.

## DISCUSSION

STING activation in tumors can promote anti-tumor immune responses, and STING agonists represent an emerging class of immunotherapy agents (13, 42, 43). However, multiple reports also indicate that STING can induce immunosuppressive tumor environments (reviewed in 44). Several pro-tumor mechanisms of STING have been described, including upregulation of programmed death ligand (44) and indoleamine 2,3-dioxygenase (45), radiation-induced recruitment of myeloid suppressor cells (46), and DMBA exposure (47). One example of immune equilibrium mediated by STING is the polarization of macrophages, where STING has been reported to induce both anti-tumor M1 as well as pro-tumor M2 subtypes (44, 48, 49).

Our lab and others have found that STING activation can also increase expression of the chemokine CCL22 in cancer cells (17–19). CCL22 is implicated in tumor immune evasion through its recruitment of regulatory T cells (25, 26) and, more recently, by promoting polarization of the M2 phenotype in tumor-associated macrophages (27). We previously showed that STING activation by cytosolic dsDNA and the STING agonist cGAM(PS)2 can upregulate *CCL22* expression in epithelial cancer cells via the STING-IRF3 axis (19). In the present study, we show that macrophages and monocytes do not themselves upregulate *CCL22* in response to STING activation but rather release unidentified factor(s) that in turn increase *CCL22* expression in a paracrine fashion in cancer cells. STING-activated myeloid cells that can induce neighboring cancer cells to upregulate CCL22 via paracrine signaling may thus contribute to immunosuppressive tumor environments by favoring the recruitment of Tregs and polarization of M2a macrophages.

Based on our prior finding that *CCL22* upregulation was downstream of the STING-IRF3 axis, we hypothesized that type I IFNs could be the relevant paracrine factor(s). However, treatment of cancer cells with exogenous IFNα or IFNβ did not, in most cases, achieve the same level of *CCL22* upregulation as conditioned media from STING-activated myeloid cells. For example, in HeLa cells, treatment with type I IFNs increased *CCL22* by 10-20 fold, while treatment with conditioned media from STING-activated macrophages increased *CCL22* by up to 8,000 fold. In MCF7 cells, conditioned media from monocytes, but not macrophages, also increased *CCL22* above levels induced by type I IFNs, although the effect was more modest.

Surprisingly, we found that *CCL22* upregulation by exogenous type I IFNs, as well as paracrine factors, remained dependent on STING, as evidenced by the lack of *CCL22* upregulation in STING knockout cells. Although STING-dependent expression of type I IFNs is well-described, a requirement for STING for a functional effect of exogenous type I IFNs has, to our knowledge, not been previously described. One possible explanation could be our finding that IFNβ itself induces activating phosphorylation of STING, which also has not been previously reported. However, other studies have indicated connections between type I IFNs and the transcriptional regulation of *cGAS* and the gene that encodes STING, *TMEM173* (50, 51).

These findings raise important questions about feedback mechanisms between type I IFNs and STING. Regarding such feedback, LINE-1 retrotransposons, which are reported to activate cGAS-STING (5), were recently shown to be upregulated in both monocytes and neutrophils exposed to IFNα (52). Many solid tumors upregulate LINE-1 expression (9, 53), making LINE-1 a possible source of CCL22 in cancer cells via cGAS-STING activation. The potential for additional LINE-1 upregulation in myeloid cells by type I IFNs may further amplify STING signaling in those cells, increasing the release of paracrine factors affecting *CCL22*.

More studies are needed to identify the paracrine factor(s) released from STING-activated myeloid cells that induce *CCL22*. Multiple reports show that a variety of factors can influence CCL22 regulation, including but not limited to IL-1, IL-4, IL-5, IFN-gamma (IFN-γ), and tumor necrosis factor alpha (TNF-α), with some factors having different effects on CCL22 based on cell type (e.g. 28, 29, 54-59). A study in 2011 by Faget et al. showed that peripheral blood mononuclear cells promoted increased secretion of CCL22 from breast cancer cells and identified roles for monocyte-derived TNF-α and IL-1β, as well as exogenous IFN-γ (60). Whether any of these factors, or combinations thereof, are among the paracrine factors released by STING-activated myeloid cells in our study remains to be determined. However, one important potential candidate, especially given the STING-dependency we observed for *CCL22* upregulation in cancer cells, could be cGAMP itself. Intercellular transfer of cGAMP has been shown to occur via several mechanisms, including cell-cell contacts such as phagocytosis and gap junctions, as well as the release and uptake of cGAMP from the extracellular space through various transport channels in the cell membrane (4, 61–65). Although these studies have largely focused on cGAMP released by tumor cells that is taken up by myeloid cells, the converse may also be true.

The findings presented here offer new insights into possible mechanisms of STING-mediated immunosuppression in cancer. Future studies will be needed to determine clinical associations between STING activation and increased intratumoral CCL22, and whether increased levels of CCL22 in this context correlate with immunosuppression. In addition, identification of the paracrine factor(s) released from STING-activated myeloid cells that can increase CCL22 in cancer cells could reveal therapeutic strategies targeting Treg recruitment and M2 polarization in solid tumors.

---

**Supplementary Figure 1.**
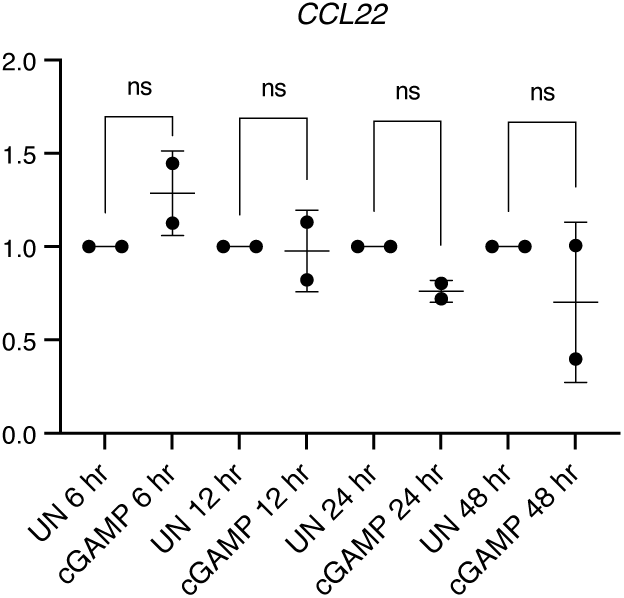
THP-1-derived macrophages were untreated (UN) or treated with cGAM(PS)2 (30 μM) for indicated times, then harvested in TRIzol for RNA purification and RTqPCR. Relative gene expression of *CCL22* is shown. Error bars represent standard deviations of independent biological replicates. Statistical significnace was determined uisng Kruskal-Wallis multiple comparisons test.

**Supplementary Figure 2.**
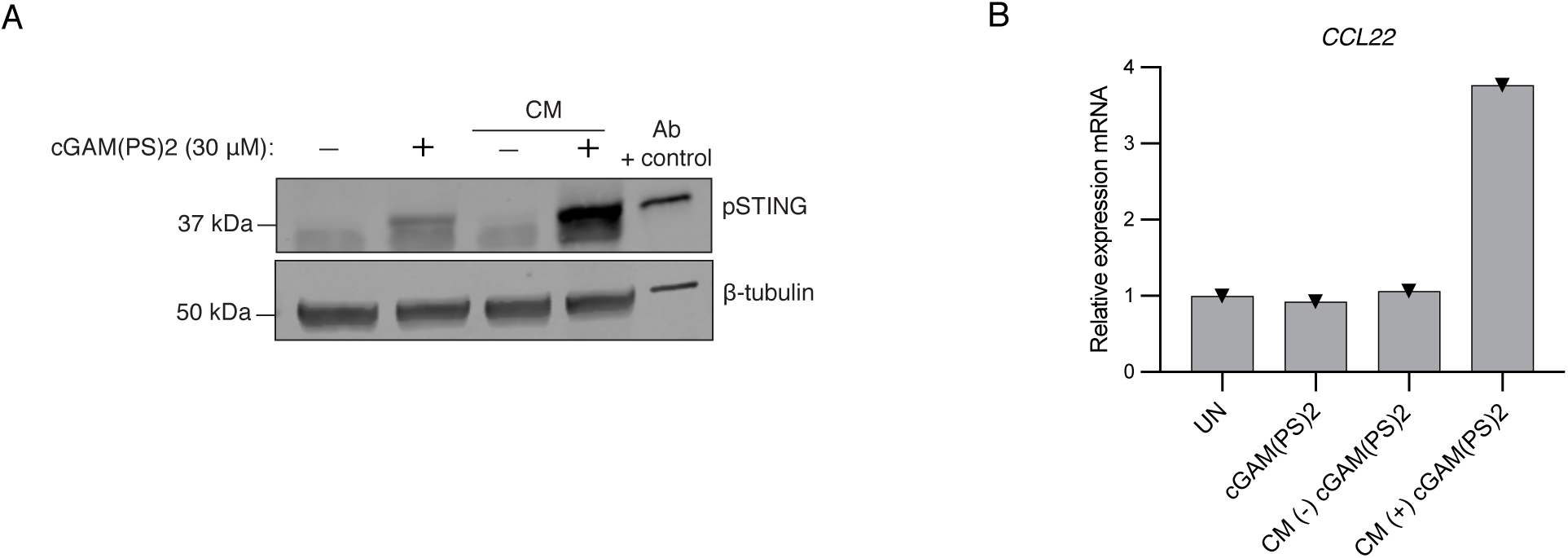
***A)*** HeLa cells (wild type) were untreated or treated for 24 hours with either cGAM(PS)2 (30 μM) or conditioned media (CM) from untreated monocytes or monocytes treated with cGAM(PS)2 (30 μM) for 24 hours. Whole cell lysates (60 μg) and antibody positive control (15 μg) were resolved with SDS-PAGE, transferred to nitrocellulose and probed for phosphorylated STING and beta-tubulin. ***B)*** HeLa ^-/-TMEM173^ (STING KO) cells were untreated or treated with either cGAM(PS)2 (30 μM) or conditioned media (CM) from untreated monocytes or monocytes treated with cGAM(PS)2 (30 μM) for 24 hours. Cells were harvested in TRIzol for RNA purification and RTqPCR to determine relative levels of *CCL22*.

**Supplementary Figure 3.**
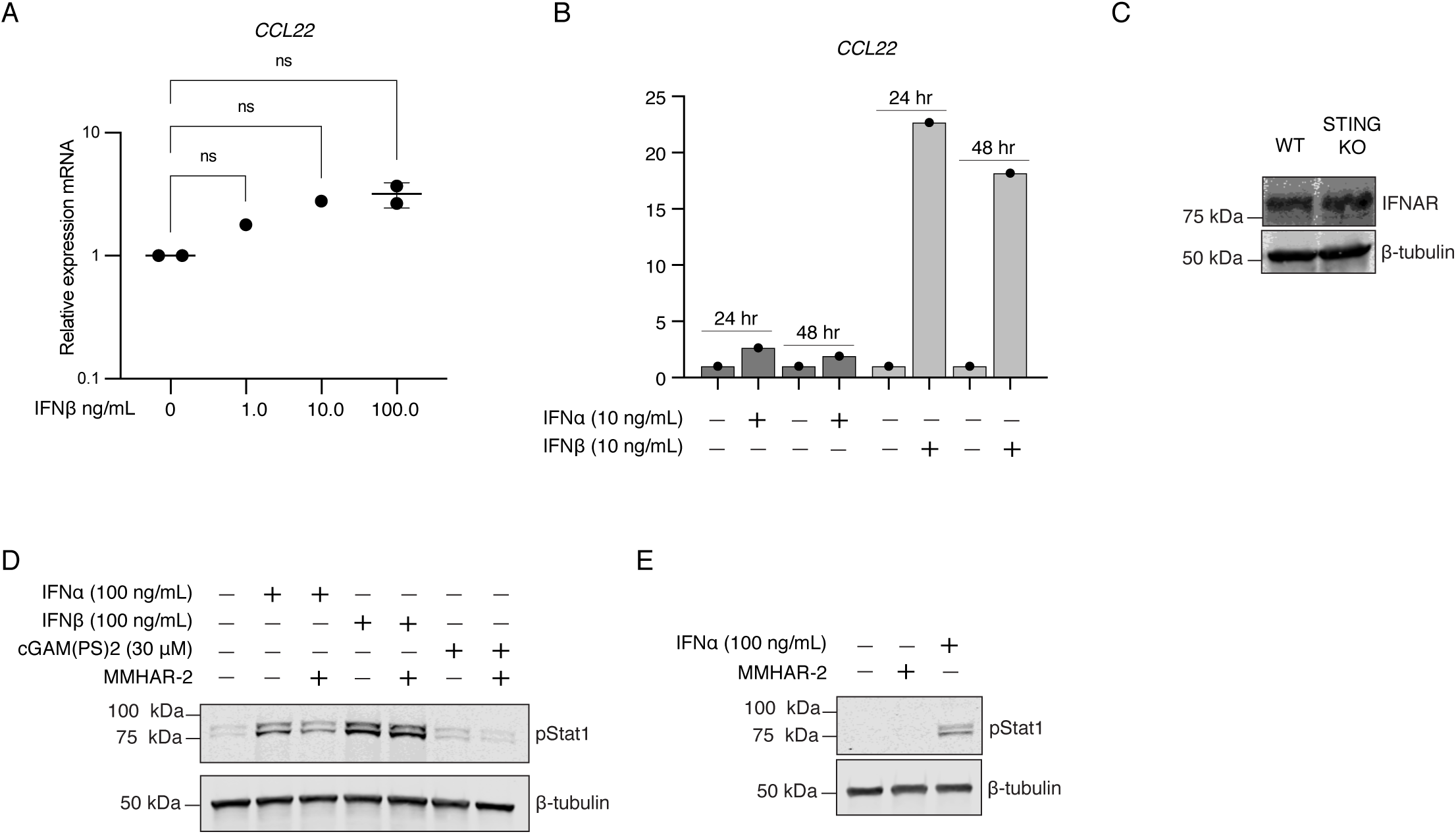
***A)*** HeLa cells were treated for 24 hours with exogenous IFNβ at indicated concentrations, then harvested in TRIzol for RNA purification and RTqPCR to assess relative fold changes in *CCL22* expression. Error bars represent standard deviations of independent biological replicates, and statistical analysis was performed with one-way ANOVA and Dunnett’s pairwise comparsions. ***B)*** MCF7 cells were treated for 24 or 48 hours with 10 ng/mL of IFNα or IFNβ, then harvested for RTqPCR as in *A*. ***C)*** Whole cell lysates (40 μg) from HeLa wild type (WT) and HeLa ^-/-TMEM173^ STING knock out (KO) cells were resolved with SDS-PAGE, transferred to nitrocellulose, and probed for IFNAR2 and beta-tubulin. ***D-E)*** HeLa cells were pretreated for 3 hours with MMHAR-2 neutralizing antibody, then indicated treatments were added and cells incubated for approximately 48 hours, at which time whole cell lysates (5 μg) were resolved with SDS-PAGE, transfered to nitrocellulose and probed with anti-pStat1 and β-tubulin.

## Notes

### Competing Interest Statement

The authors have declared no competing interest.

### Summary of Updates

Title change, clarifications in text, revised discussion.

